# Transcriptomic analyses revealed the effect of *Funneliformis mossea*e on differentially expressed genes in *Fusarium oxysporum*

**DOI:** 10.1101/2020.05.28.120873

**Authors:** Xue-Qi Zhang, Li Bai, Na Guo, Bai-Yan Cai

## Abstract

Soybean root rot is a typical soil-borne disease that severely affects the yield of soybean, and *F. mosseae*, the dominant strain of AMF in continuous cropping of soybean. The aim of this study was to providing an experimental basis for the study of the molecular mechanism underlying the alleviation of the obstacles associated with the continuous cropping of soybean by AMF. In this study, *F. mosseae* was inoculated in soil planted with soybean infected with *F. oxysporum*. The results showed that the incidence of soybean root rot was significantly reduced after inoculation with *F. mosseae*. The significantly upregulated genes encoded the ABC transporter, ATP-binding/permease protein and the ABC transporter, ATP-binding protein. The significantly downregulated genes encoded chitin-binding domain proteins; key enzymes involved in metabolic pathways such as glycolysis, including class II fructose-bisphosphate aldolase and NAD-dependent glyceraldehyde-3-phosphate dehydrogenase, glycoside hydrolase family 61 protein, which hydrolyse cellulose and hemicellulose; actin and other major components of the cytoskeleton. The DEGs were enriched in antigen processing and presentation, carbon fixation in photosynthetic organisms, glycolysis/gluconeogenesis, the MAPK signalling pathway, protein processing in the endoplasmic reticulum and RNA degradation. Inoculation with *F. mosseae* could promote the growth and development of soybean and improve disease resistance. This study provides an experimental basis for further research on the molecular mechanism underlying the alleviation of challenges associated with the continuous cropping of soybean by AMF.

## Introduction

*Fusarium* root rot of soybean is a kind of soil fungal disease that is widely distributed, harmful and difficult to control. This disease occurs all over the world[1]. The annual decline of soybean yield caused by root rot can reach an average of 1-30% and up to nearly 50%.

Crop root rot caused by *Fusarium oxysporum* (*F. oxysporum*) is a typical destructive soil-borne disease. This pathogen has many specialized types and a wide host range, causing more than a 100 diseases in different plants, such as melons, solanaceae, legumes and flowers [2]. This pathogen infects plant roots, causing vascular diseases. *F. oxysporum* exhibits strong virulence and high infection and mortality rates. At the onset of infection, the root begins to exhibit discolouration from the apex; the lower part of the main root first exhibits brown spots, which gradually expand, and the epidermis and cortex become black and rotten. In severe cases, the lower part of the main root rots, the leaves gradually turn yellow from bottom to top, the plant size deceases and the number of pods decreases [3–4]. *F. oxysporum* poisons the root systems of host plants by secreting fusarin and cell wall degrading enzymes [5]. Currently, there is no specific method to prevent and control *F. oxysporum* infection in agricultural production, and there has been little research on the pathogenesis of *F. oxysporum*.

*Funneliformis mosseae*, as a dominant strain of arbuscular mycorrhizal fungi (AMF), exists in the rhizosphere and plant tissues and plays a key role in plant evolution and nutrition as an obligate symbiont [6]. Studies have shown that AMF can improve plant stress resistance, such as saline-alkali resistance, heavy-metal resistance, drought resistance [7–8], and disease resistance (resistance to root rot, Verticillium wilt, blight, etc.)[9–12]. Guo et al. found that after inoculation with *F. mosseae*, the symbiotic mycorrhizal network formed between *F. mosseae* and donor plants increased the activities of phenylalanine ammonialyase, polyphenol oxidase, and peroxidase in tobacco plants [13]. However, there are few reports on the mechanism by which *F. mosseae* inhibits the pathogenicity of *F. oxysporum*, the main pathogen associated with root rot.

In this study, *F. mosseae*, the dominant strain of AMF in continuous cropping of soybean, was inoculated into the soil of soybean plants infected with *F. oxysporum*. Illumina HiSeqTM sequencing was used to investigate the differences in gene transcription in *F. oxysporum* at the transcriptome level after inoculation with *F. mosseae*, thus providing an experimental basis for the study of the molecular mechanism underlying the alleviation of the obstacles associated with the continuous cropping of soybean by AMF.

## Results

### Incidence of root rot

The root systems of F-vs.-AF were observed. The appearance of root rot is shown in Figure 1. In Figure 1A, the root system of plant inoculated with *F. oxysporum* (F) showed blackening, while this phenomenon was less obvious in the roots of AF. The lesion at the inoculation area (Figure 1B) was large. As shown in Figure 2, the incidence of root rot disease became increasingly severe with the growth and development of soybean. Compared with AF, the incidence of root disease in soybean group F increased significantly. AF and F showed the same trend at the initial stage. With the growth of soybean, the incidence of AF gradually slowed down and eventually approached the level of the control group (CK). The highest incidence in group F was almost 90%. The root rot in soybean inoculated with *F. mosseae* was significantly reduced, indicating that *F. mosseae* could significantly alleviate the incidence of soybean root rot.

**Figure 1.**
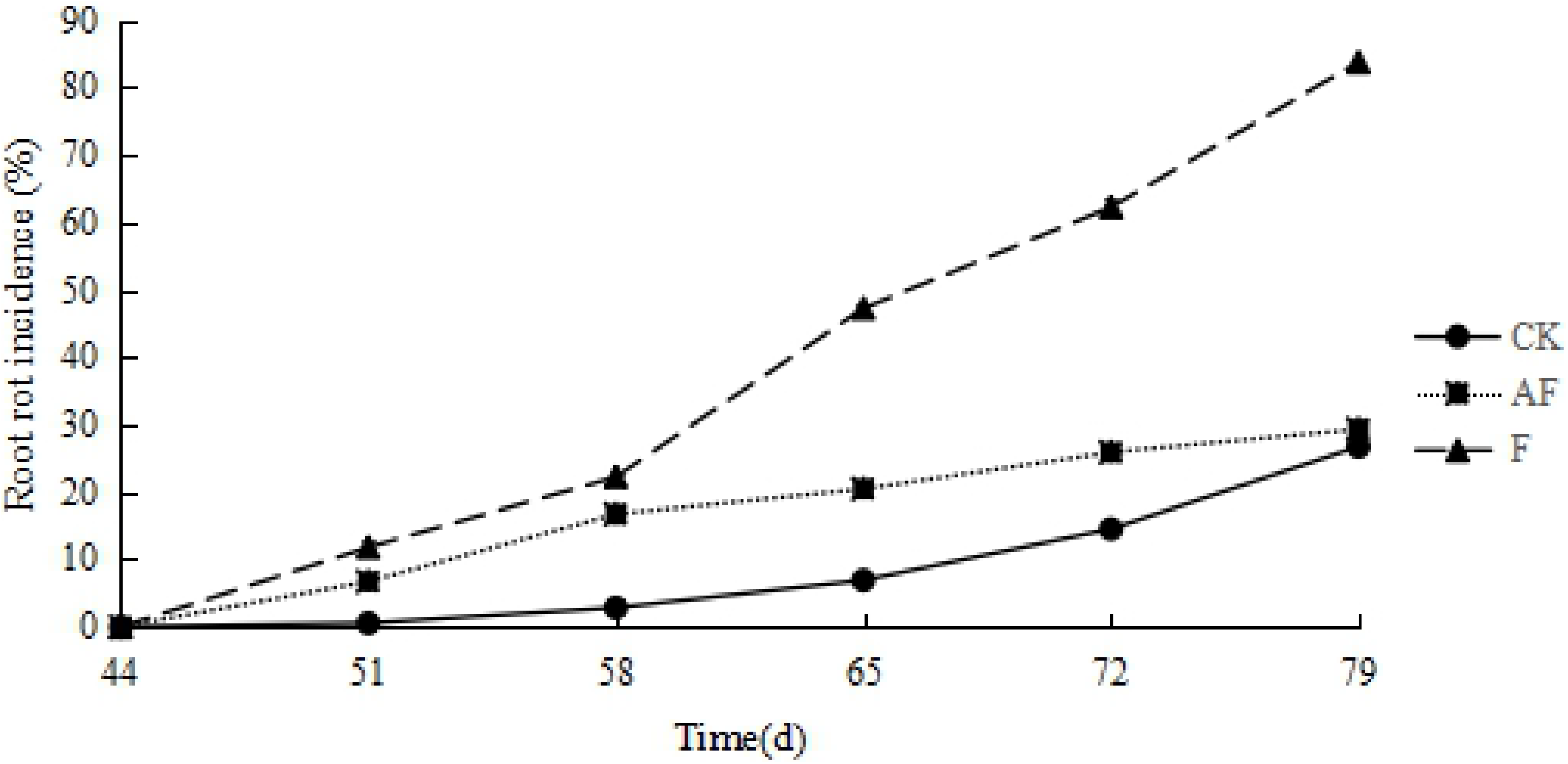
Soybean root rot. (A) The root system of plant inoculated with *F. oxysporum* shows roots of the group that was only inoculated with the pathogenic fungus *F. oxysporum* (left, F) and simultaneous inoculation of *F. mosseae* and the pathogenic fungus *F. oxysporum* (right, AF). (B) Prominent in the red frame is the disease spot of root system inoculated with *F. oxysporum*.

**Figure 2.**
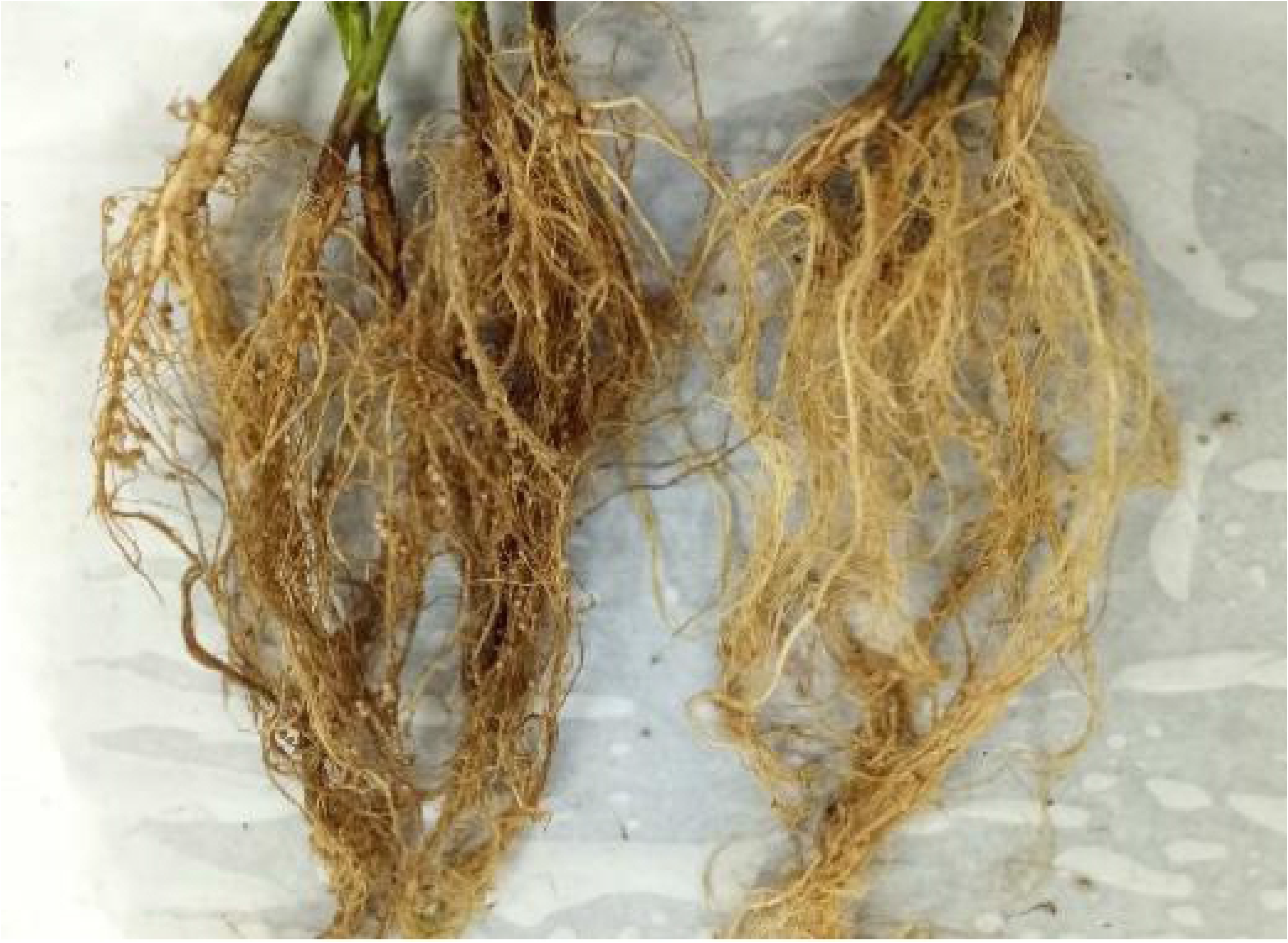
Root rot incidence. CK, control group; AF, soybean planted in continuously cropped soil inoculated at 44 d after sowing and evenly mixed with F. mosseae; F, soybean planted in continuously cropped soil inoculated with F. oxysporum at 44 d after sowing. The numbers 44, 51, 58, 65, 72 and 79 represent the days after sowing. The x-axis means different stages. The y-axis represents the soybean root rot incidence.

### Soybean growth indexes

After soybean ripening, soybean grains were randomly picked for determination of 100-grain weight, crude fat content and protein content, as shown in Figure 3. Root rot caused by *F. oxysporum* affected the fat, protein content and the 100-grain weight of soybean severely. However, inoculation with *F. mosseae* could significantly alleviate the growth depression caused by root rot and promote plant nutrition and yield.

**Figure 3.**
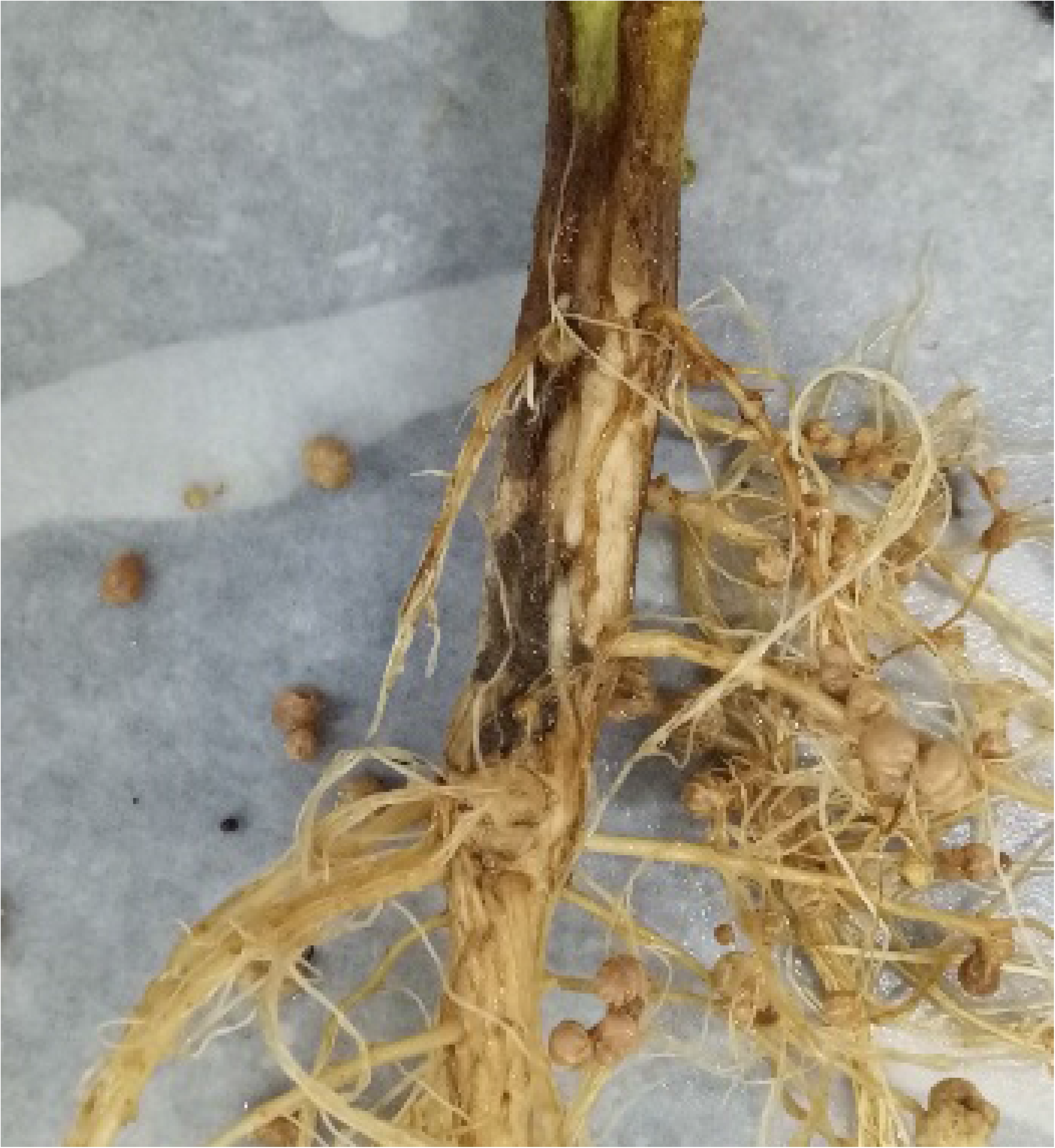
Soybean growth indexes. The x-axis means different treatment groups, CK, control group; AF, soybean planted in continuously cropped soil inoculated at 44 d after sowing and evenly mixed with *F. mosseae*; F, soybean planted in continuously cropped soil inoculated with *F. oxysporum* at 44 d after sowing. (A) The y-axis means soybean fat content. (B) The y-axis means soybean protein content. (C) The y-axis means soybean 100-grain weight.

The correlation coefficients between the incidence of root rot and soybean quality indicators are shown in Table 1, which shows that the incidence of soybean root rot is negatively correlated with various quality indicators (P < 0.01), and the increase in disease incidence has the most serious impact on the soybean fat content, which decreases upon infection. In addition, the protein content and fat content of soybean were positively correlated with the 100-grain weight (P < 0.01), and the fat content had a strong impact on the 100-grain weight of soybean.

**Table 1.** Coefficients of correlation between the incidence of root rot and protein content, 100-grain weight and fat content.

| Index | Incidence | Protein content | 100-grain weight | Fat content |
| --- | --- | --- | --- | --- |
| Incidence | 1 | -.870** | -.972** | -.982** |
| Protein content |  | 1 | .920** | .899** |
| 100-grain weight |  |  | 1 | .977** |
| Fat content |  |  |  | 1 |

Note: \*\* Significant difference, $P < 0.01$

### Transcriptomic Analysis of *F. oxysporum* in Soybean Roots

#### Gene expression analysis

The high-quality clean reads obtained were compared with known genomes. Finally, 8398, 5265 and 5110 genes were detected in the CK, AF and F treatments, and the number of genes detected in each group is shown in Table 2.

**Table 2.**
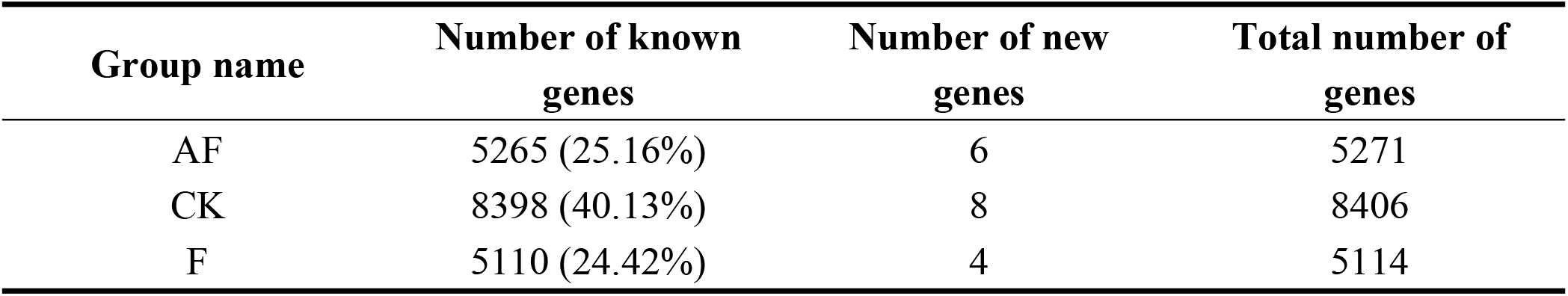
Number of genes detected in each treatment group.

| Group name | Number of known genes | Number of new genes | Total number of genes |
| --- | --- | --- | --- |
| AF | 5265 (25.16%) | 6 | 5271 |
| CK | 8398 (40.13%) | 8 | 8406 |
| F | 5110 (24.42%) | 4 | 5114 |

The comparisons of gene expression levels between groups are shown in the box plot in Figure 4. The y-axis represents gene expression levels, and each box represents a sample group. The black line in the middle of the box represents the median, and the upper and lower sides of the box are quartiles. Based on the box plot, the number of genes detected in each treatment group was different.

**Figure 4.**
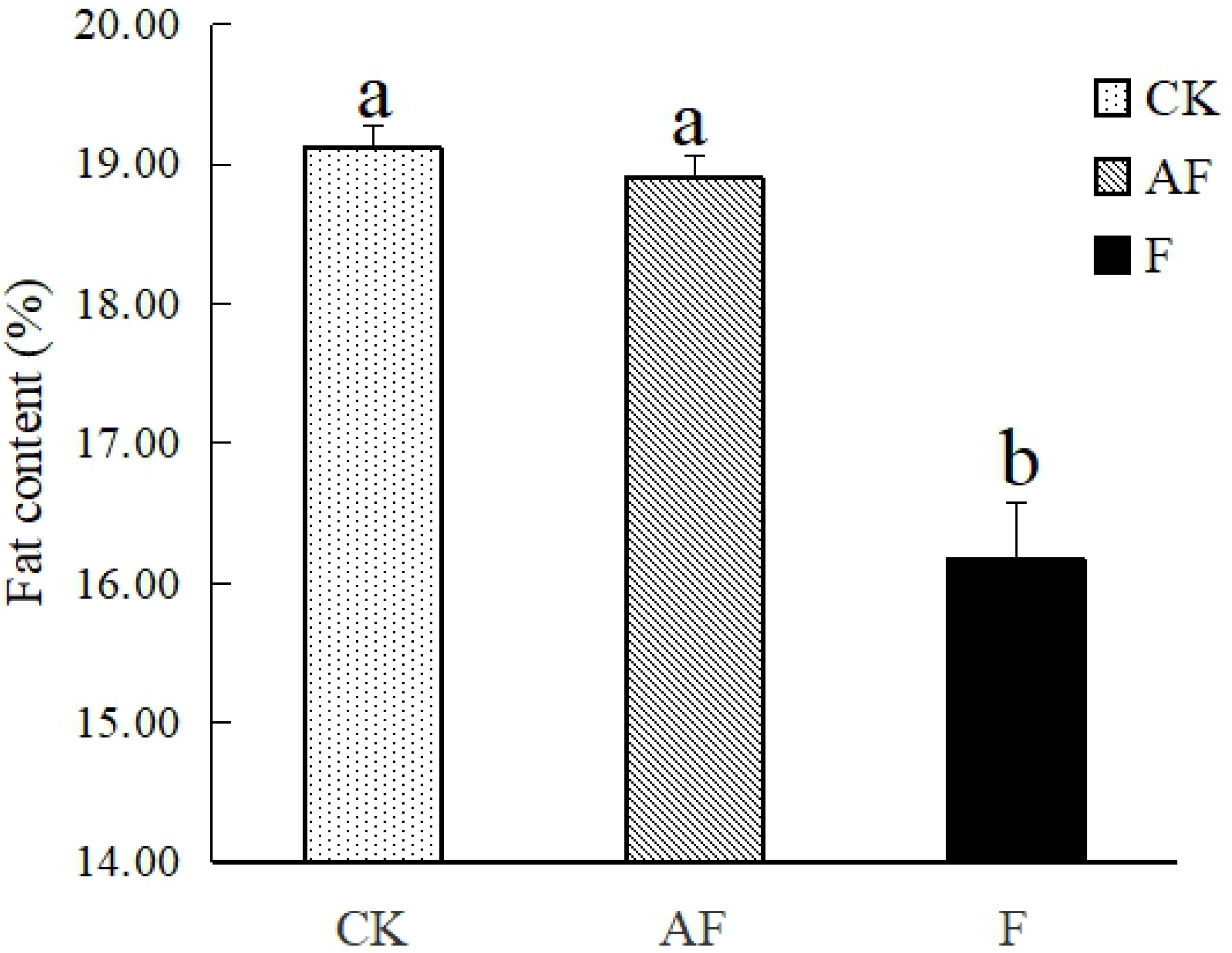
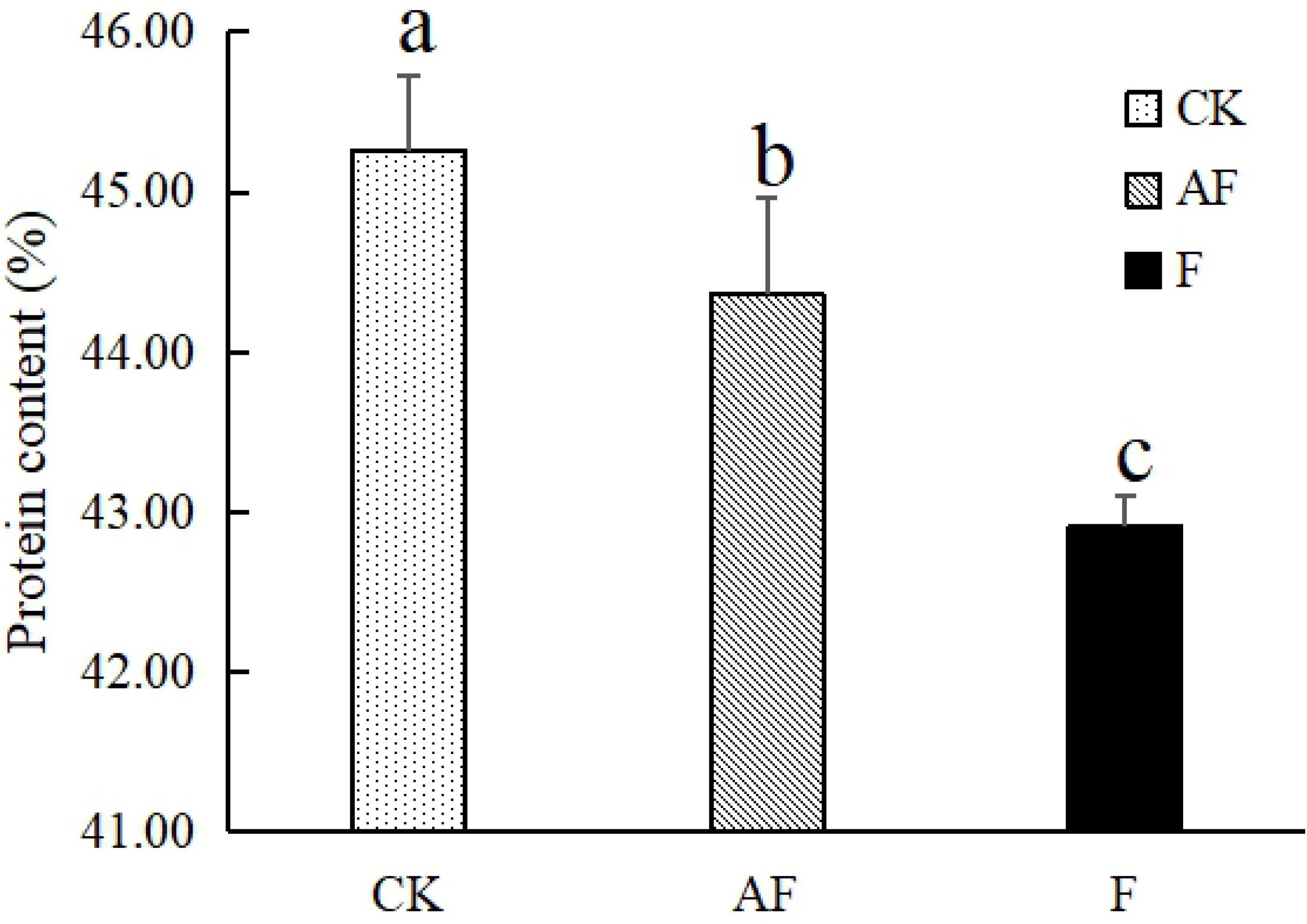
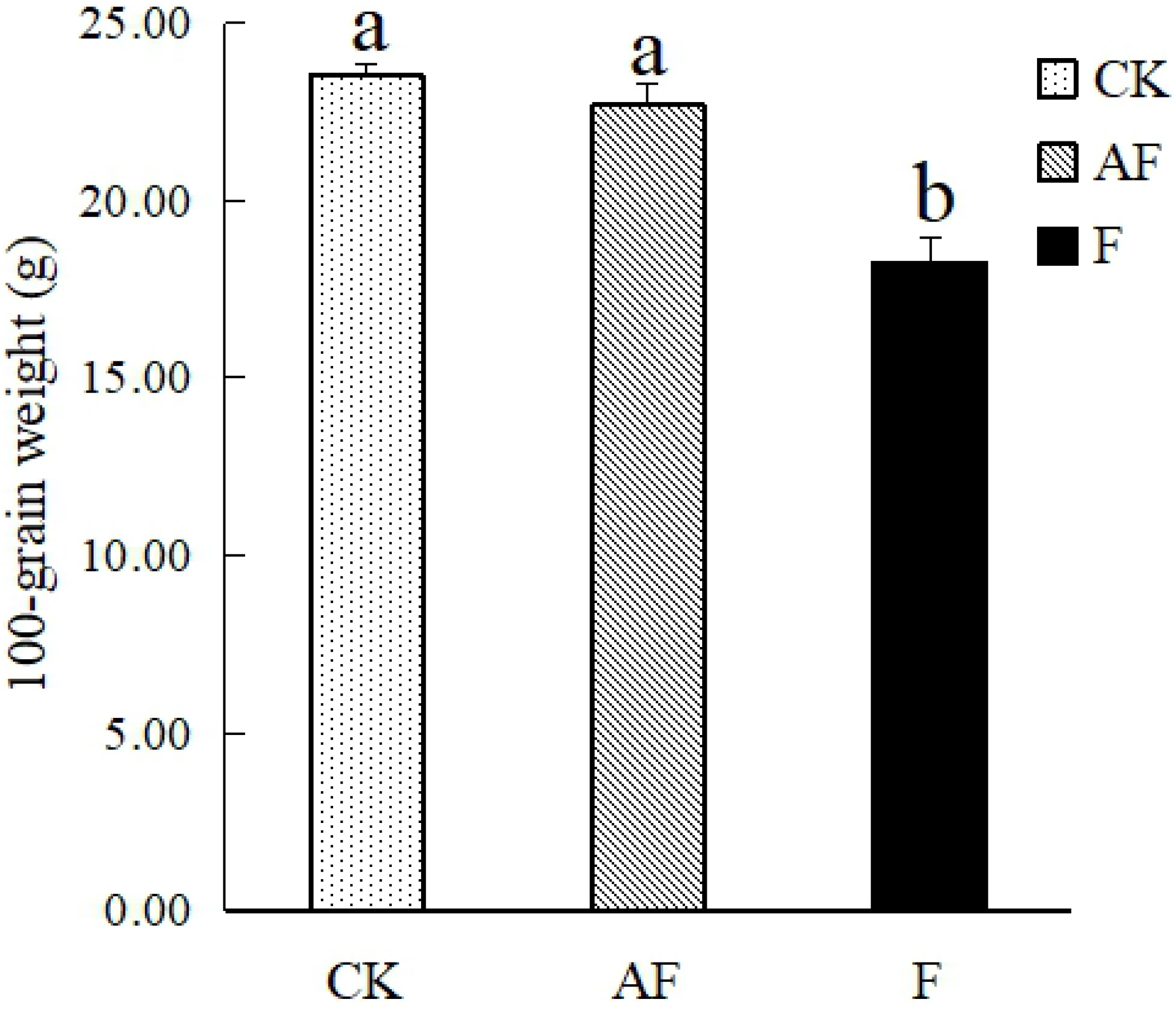
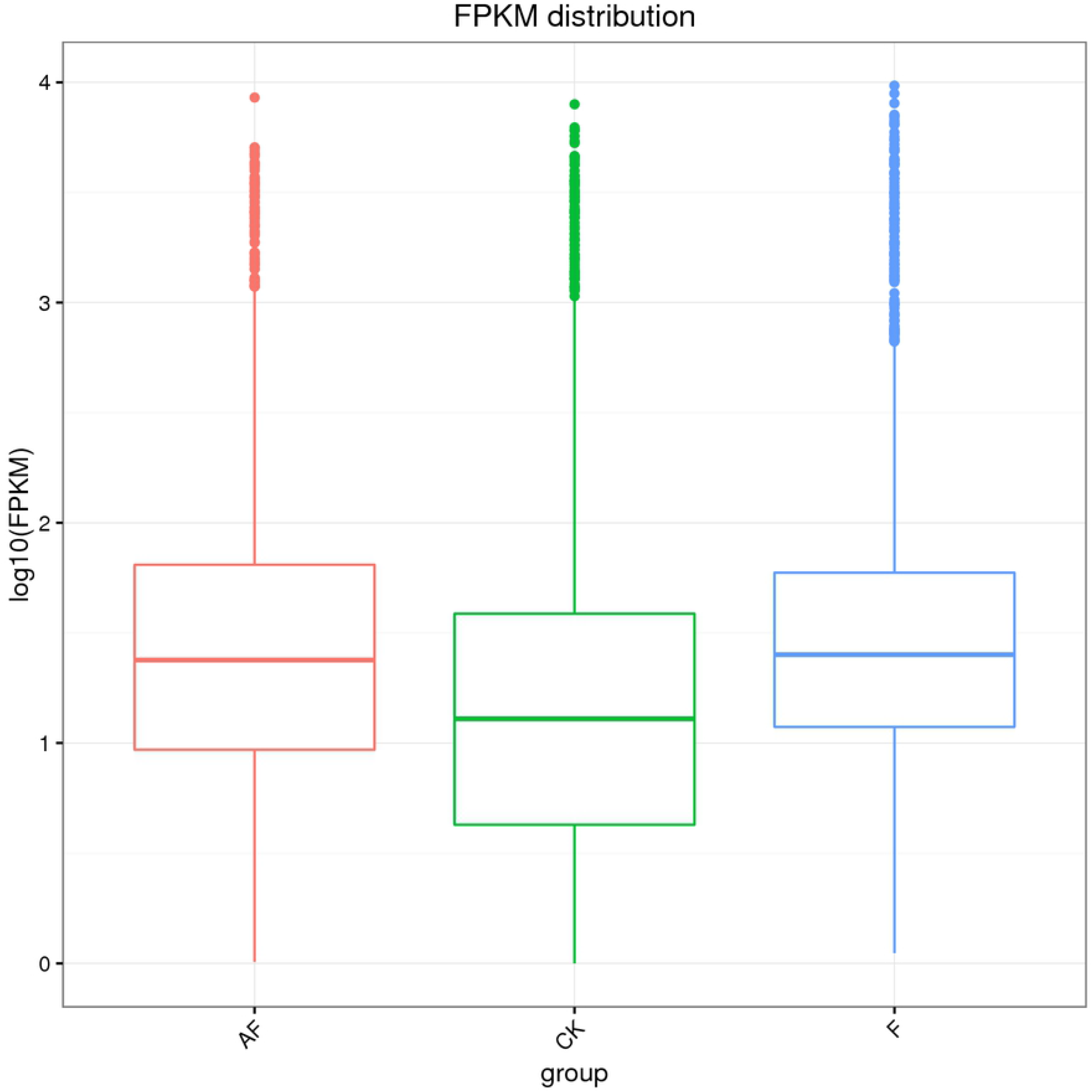
Box plot for comparing gene expression levels between groups. The vertical coordinate in the figure is the gene expression quantity, and each box represents a sample group. The line in the middle of the box represents the median, and the upper and lower sides of the box are the upper and lower quartiles respectively.

#### Sample relationship

Principal component analysis (PCA) was used to determine the effect of inoculation with *F. oxysporum* on soybean and the effect of *F. mosseae* on *F. oxysporum* in susceptible soybean roots (Figure 5). PC1 accounted for 58.9% of the total variance of all variables (expression levels of all the genes). PC1 and PC2 accounted for 73.7% of the total variance. Figure 5 shows that in this processing range, the single pathogen has a strong impact on gene expression in the samples. After inoculation with *F. mosseae*, the gene expression levels of soybean in the AF group were slightly different from those in the CK group, which indicated that *F. mosseae* could inhibit the pathogenicity of *F. oxysporum*.

**Figure 5.**
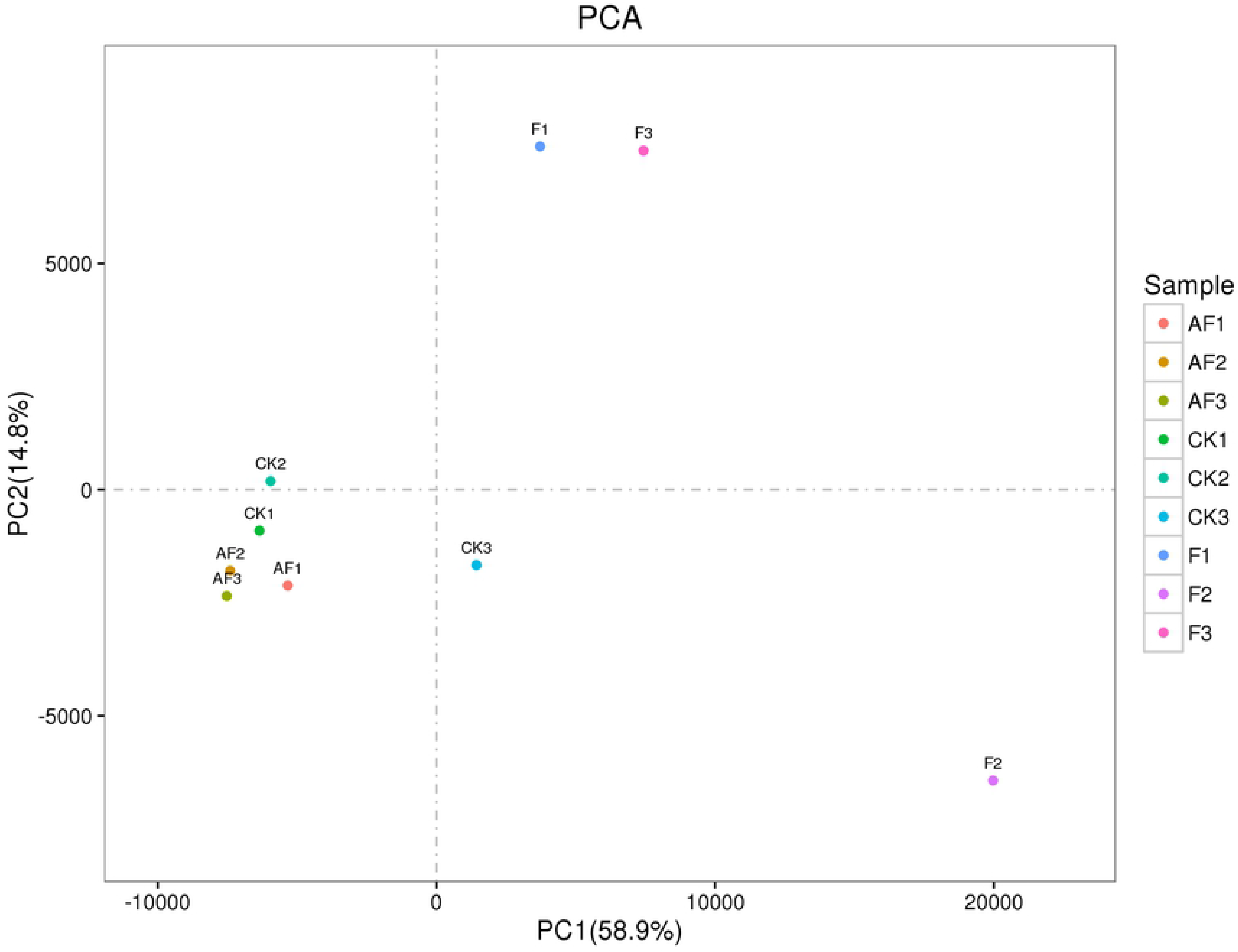
PCA. According to the value of each sample in the first principal component (PC1) and the second principal component (PC2), two-dimensional coordinate map is made. The values in brackets on the axis labels represent the percentage of the variance of the population explained by the principal component. PC1 can explain 55% of the total variance of all variables (expression of all genes), PC1 and PC2 can explain 91.7% of the total variance.

### Analysis of differentially expressed genes (DEGs) of *F. oxysporum*

The difference in gene expression among the three treatment groups was analysed by edgeR software. In F-vs.-AF, the number of upregulated was 56 and downregulated genes is 35. Hierarchical clustering of the relationships between samples and genes was carried out based on gene expression. The clustering results are presented using a heatmap, as shown in Figure 6, where red indicates high expression levels, and blue indicates low expression levels.

**Figure 6.**
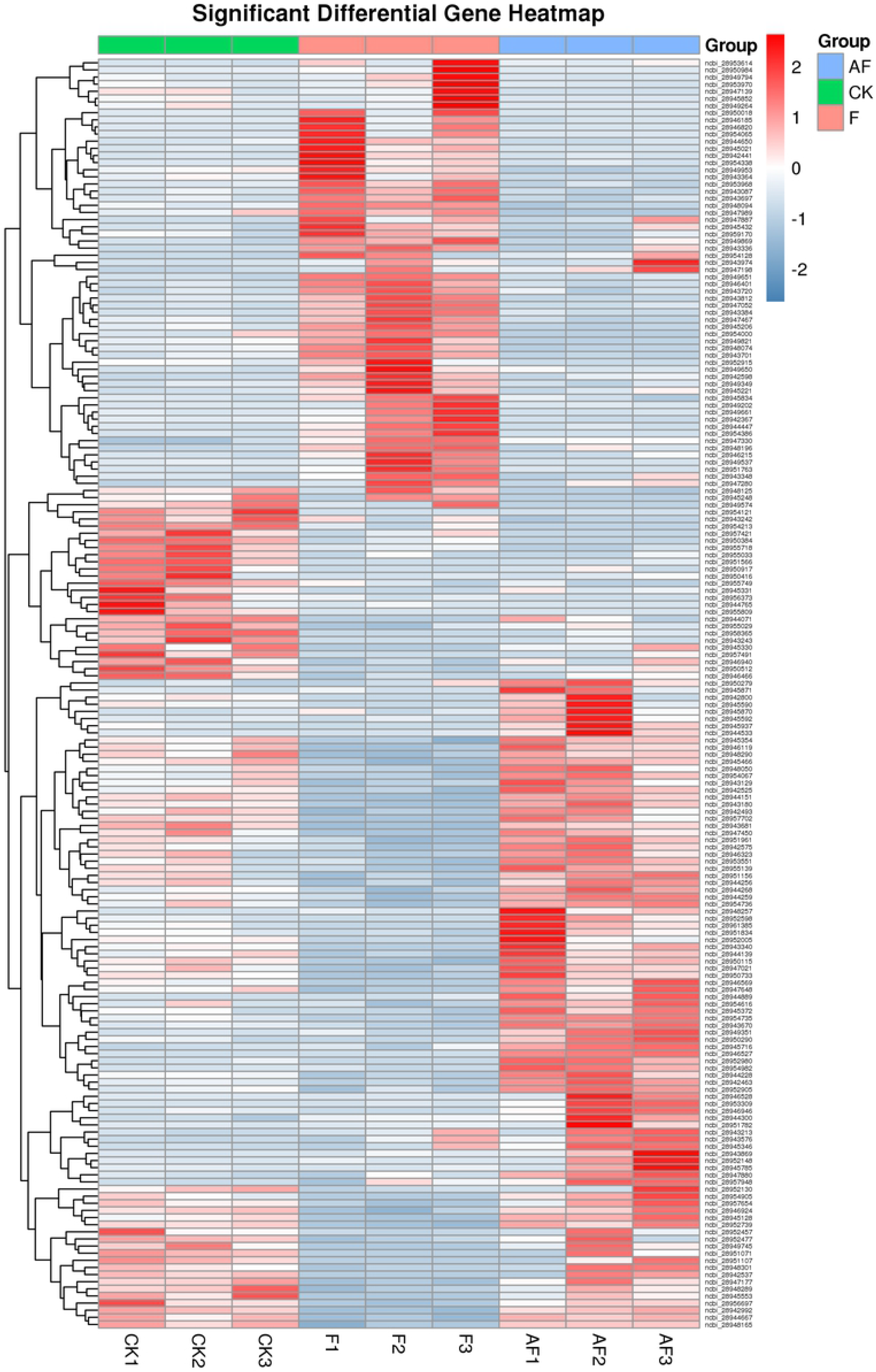
Clustering of differential expression patterns between groups. Based on the amount of gene expression, the hierarchical clustering of the relationship between samples and genes is carried out, and the results of clustering are presented by using heat map. Take 2 as the base to calculate the logarithm value of gene expression of each sample, and carry out hierarchical cluster analysis for different samples and genes. Each column in the figure represents a sample, each row represents a gene, and the expression amount of gene in different samples is expressed in different colors. The redder the color, the higher the expression, and the bluer the color, the lower the expression.

As shown in Figure 6, after inoculation with *F. mosseae*, the significantly upregulated genes were ABC transporter, ATP-binding/permease protein-encoding genes and ABC transporter, ATP-binding protein-encoding genes. Significantly downregulated genes were chitin-binding domain protein-encoding genes, genes encoding key enzymes involved in metabolic pathways such as glycolysis, including class II fructose-bisphosphate aldolase and NAD-dependent glyceraldehyde-3-phosphate dehydrogenase, glycoside hydrolase family 61 protein, which are involved in the hydrolysis of cellulose and hemicellulose, and genes encoding actin and other major components of the cytoskeleton. Chitin-binding domain proteins are a class of chitin-specific binding proteins that usually contain one or more chitin-binding domains[14]. These proteins can participate in the recognition or binding of chitin and chitin subunits and may play a special role in the interactions between plants and pathogens.

### Gene Ontology (GO) analysis

GO was used to classify DEGs after inoculation with *F. mosseae*. Biological process, molecular function and cellular component terms were annotated and classified. The enrichment results are shown in Figure 7.

**Figure 7.**
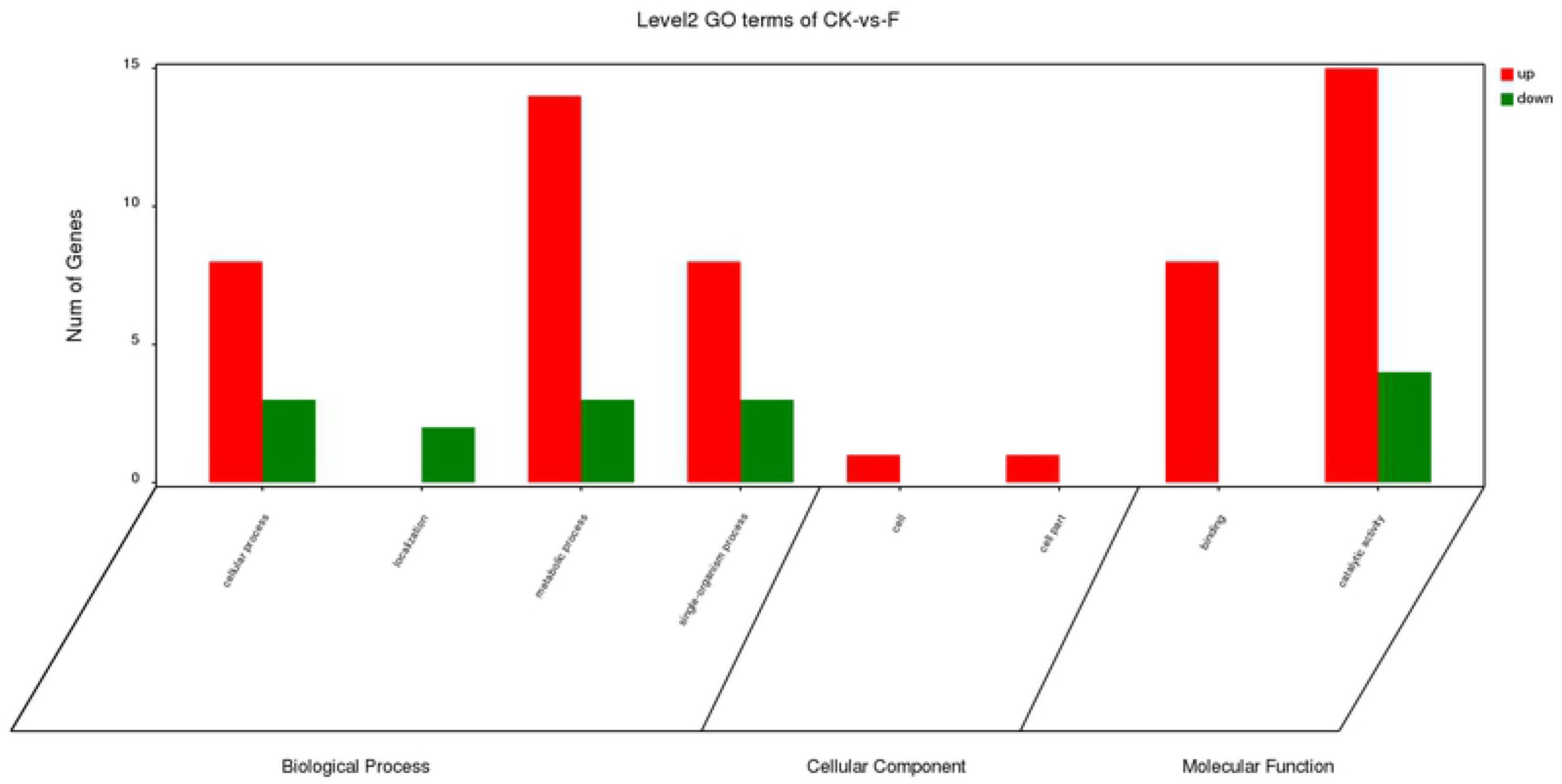
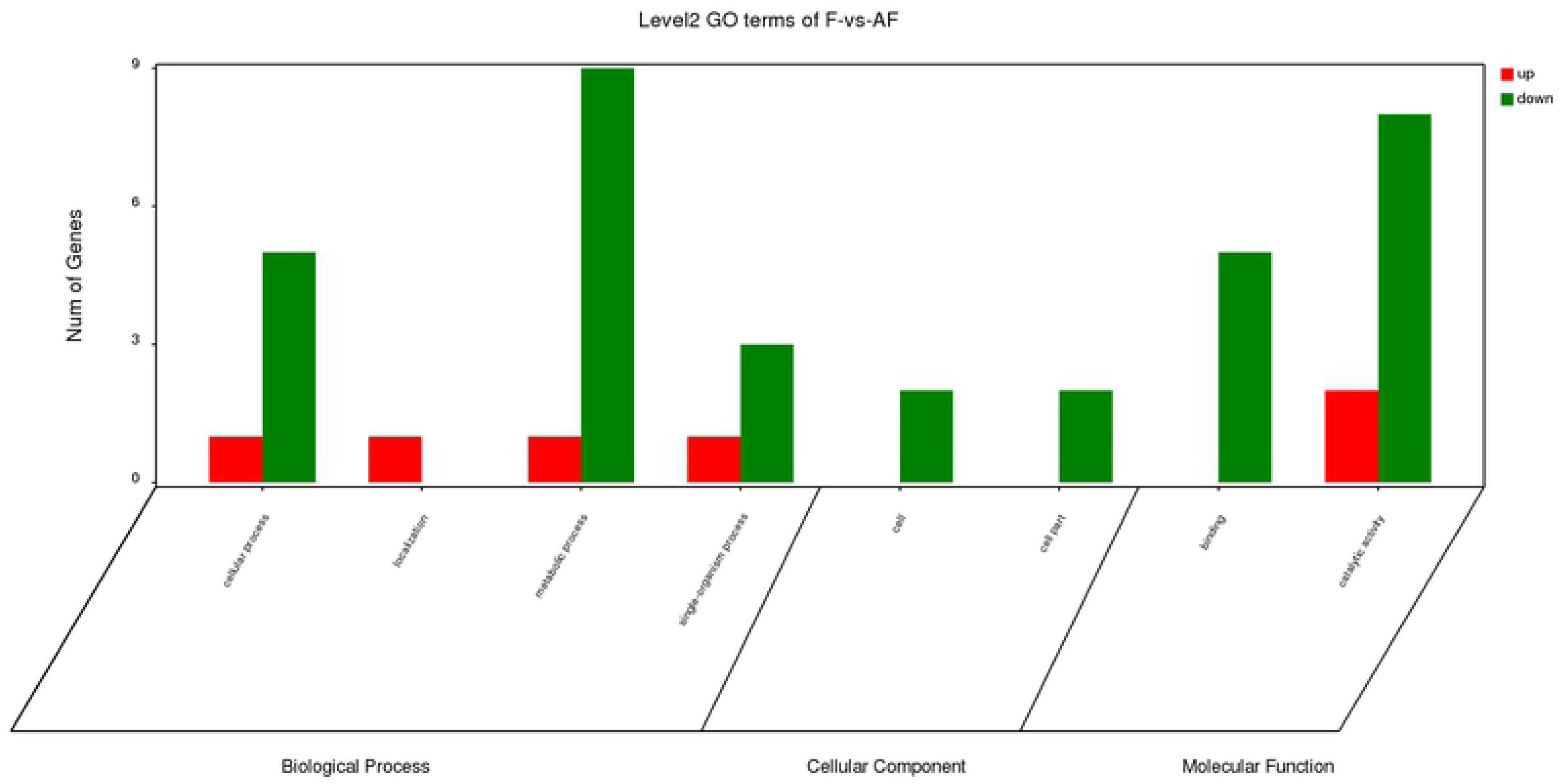
GO enrichment analysis of DEGs. DEGs in each group were sorted into three categories. Green bars indicate downregulated genes, and red bars indicate upregulated genes. The x-axis indicates different GO terms. The y-axis represents the number of genes in the indicated categories. (A) CK-vs.-F comparison; (B) F-vs.-AF comparison.

Figure 7A shows that the DEGs in the CK -vs.- F group were enriched in 8 GO terms (Table S1). In the biological process group, most of the DEGs are involved in metabolic processes(14 upregulated genes, 3 downregulated genes), cellular processes(8 upregulated genes, 3 downregulated genes) and the single-organism processes(8 upregulated genes, 3 downregulated genes). In the molecular function group, most of the DEGs are involved in catalytic activity(15 upregulated genes, 4 downregulated genes) and binding(8 upregulated genes). Figure 7B shows that the DEGs in the F -vs.- AF group were also enriched in 8 GO terms(Table S2). In the biological process group, most of the DEGs are involved in metabolic processes(1 upregulated genes, 9 downregulated genes) and cellular processes(1 upregulated genes, 5 downregulated genes). In the molecular function group, most of the DEGs are involved in catalytic activity(2 upregulated genes, 8 downregulated genes) and binding(0 upregulated genes, 5 downregulated genes).

### Kyoto Encyclopedia of Genes and Genomes (KEGG) pathway analysis

In the CK -vs.- F group, after inoculation with *F. oxysporum*, the top 20 pathways were enriched(Table S3). Figure 8A shows that many DEGs related to biochemical metabolism participate in carbon metabolism(12 DEGs), the pentose phosphate pathway(5 DEGs), glycolysis/gluconeogenesis(6 DEGs), microbial metabolism in diverse environments(12 DEGs), fructose and mannose metabolism(3 DEGs). Most of the DEGs related to biosynthesis are involved in the biosynthesis of amino acids, biosynthesis of secondary metabolites, and biosynthesis of antibiotics. Some genes are involved in signal transduction pathways, such as the oestrogen signalling pathway and NOD-like receptor signalling pathway. Some DEGs related to susceptibility and autoimmunity are involved in plant-pathogen interactions, antigen processing and presentation.

**Figure 8.**
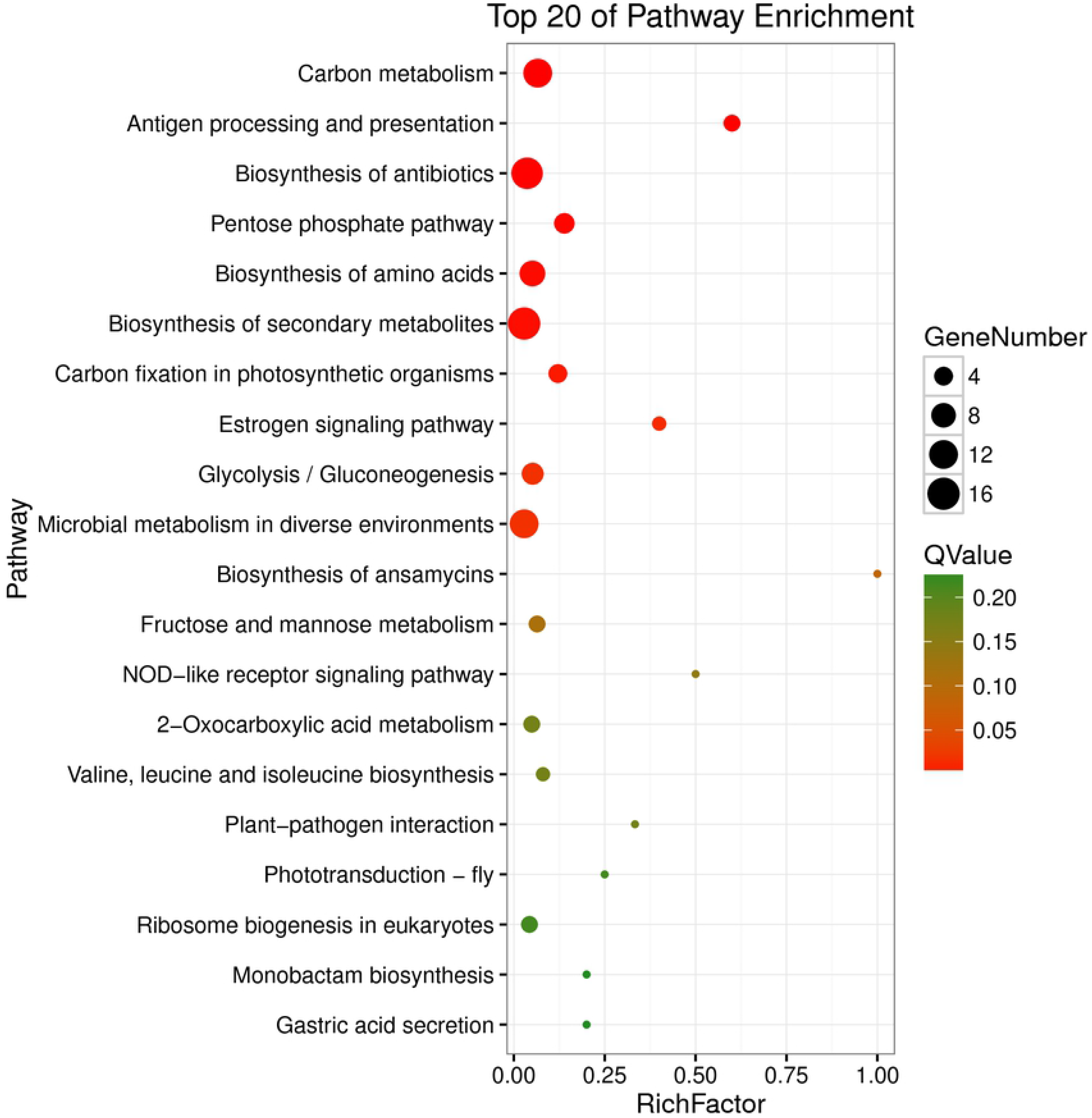
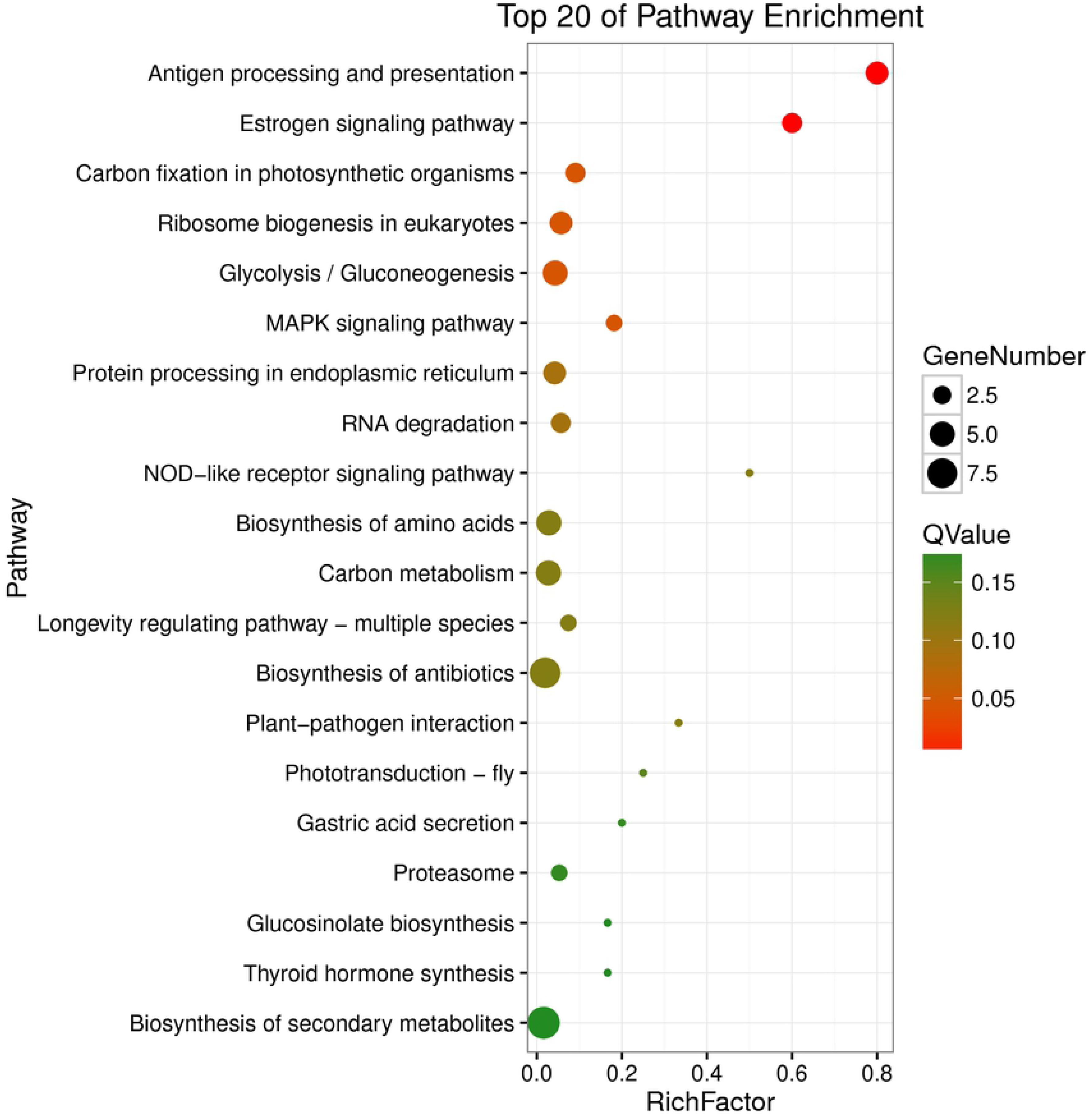
Intergroup KEGG enrichment bubble map. The Q value of the red bubble diameter is lower than that of the green bubble diameter. The bubble size is proportional to gene number. The x-axis shows the top 20 pathways in which the differentially genes expressed, and the y-axis represents rich factor. (A) CK-vs.-F comparison; (B) F-vs.-AF comparison.

In the F -vs.- AF group, the top 20 pathways were significantly enriched after inoculation with *F. mosseae*, as shown in Figure 8B. The three most genes enriched pathways are Metabolic pathways(12 DEGs), Biosynthesis of secondary metabolites(9 DEGs) and Biosynthesis of antibiotics(8 DEGs). Some of the same pathways that were enriched by DEGs in the CK -vs.- F group, such as 4 DEGs in antigen processing and presentation, 3 DEGs involed in carbon fixation in photosynthetic organisms, and 5 DEGs involed in the glycolysis/gluconeogenesis. 2 DEGs are involved in the MAPK signalling pathway, which is related to fungal virulence(Table S4). There also many DEGs were enriched in other pathways, such as endoplasmic reticulum and RNA degradation after inoculation with *F. mosseae*.

## Discussion

*F. oxysporum* is the main pathogen of soybean root rot. Previous studies have confirmed that the arbuscular mycorrhizal fungus *F. mosseae* can alleviate soybean root rot. Liu et al. found that *F. mosseae* could colonize and associate with the roots of *Amorpha fruticosa* and induce the expression of various genes during the symbiosis process[15]. Ji Weiguang et al. found that the level of *F. oxysporum* DNA in the roots and rhizospheric soil samples of soybean plants inoculated with *F. mosseae* decreased significantly[16]. To revealed the effect of *Funneliformis mosseae* on DEGs in *Fusarium oxysporum*, sterilized soil was inoculated with *F. oxysporum* alone to simulate the symptoms of root rot. The time gradient sampling method was used to calculate and observe the incidence of soybean root rot, so that the root system was sampled at the high-incidence period of soybean root rot. At the same time, determination of the related indexes of disease resistance and transcriptome sequencing were carried out. Through statistical observation of incidence and growth, it was found that *F. mosseae* could promote the growth and development of soybean. Under the same condition of root rot infection, *F. mosseae* could alleviate the plant yield reduction and increase the shoot biomass of soybean. Moreover, the fat and protein content of the treatment group inoculated with *F. mosseae* was significantly higher than that in F group when soybean riped. This result is consistent with previous research results from our laboratory and the results of Li et al.[17].

Plant cell wall is the first barrier to *F. oxysporum* infection. Cellulose and pectin are the main components of the plant cell wall. Cellulose is a macromolecular polysaccharide composed of β-1,4-glycoside bonds or β-1,3-glycoside bonds. *F. oxysporum* can secrete three cell wall-degrading enzymes, namely, pectinase, cellulase and β-glucosidase, to degrade the cell walls of plants. When the cell wall is degraded, pectin blocks the vessels of host plants, prevents the host plants from absorbing water and causes plant wilting, leading to plant death [18]. In addition to these three cell wall-degrading enzymes, *F. oxysporum* also harbours genes encoding endo-polygalacturonase, exogenous galacturonase, pectic acid endolysase and xylanase. Jonkers et al. found that knocking out any of the genes encoding such proteins by gene knockout technology would lead to apical spore formation, leading to complete or partial loss of the pathogenicity of *Fusarium* towards the host[19]. Actin aggregates to form pseudopods and filaments, increasing the adhesion of *F. oxysporum* cells to host soybeans for colonization. Then, ABC transporters are mainly responsible for the transport of toxins in host plant cells [5]. In this experiment, the actin gene of the AF group was downregulated after inoculation with *F. mosseae*, indicating that *F. mosseae* played a role in weakening the colonization by pathogenic bacteria of the host soybean. Chitin is ubiquitous in the cell walls of fungi and is mainly used to support the skeleton and protect the body. The downregulation of the chitin-binding domain protein-encoding gene in the AF group indicated that *F. mosseae* could alleviate root rot disease by destroying the cell wall of the pathogenic fungus.

KEGG analysis showed that the main enriched metabolic pathways include the MAPK signalling pathway, which plays a key role in regulating genes encoding chitin, peroxidase, beauvericin and fusaric acid. HSP70, the gene encoding MAPK, was significantly upregulated in group F, suggesting that *F. oxysporum* needed to activate the MAPK signalling pathway in vivo to induce the expression of virulence-related genes and increase toxin levels or enhance its tolerance to host immunity to infect soybean[20]. Antigen processing and presentation pathways have been shown to be associated with immune responses, transport, pathogenesis, secretion and phagocytosis. Heat shock 70 kDa protein (HSP70) is involved in antigen processing, and the presentation pathway is a highly conserved polypeptide that can be repaired by degenerative proteins to aid the physiological folding and stretching of newly synthesized polypeptide bonds, correct the misfolding of polypeptide chains, restore the functions and structures of cells, act as a ‘molecular chaperone’, support immune processes, and participate in apoptosis [21](Zhang et al., 2004). Another protein involved in the antigen processing and presentation pathway, molecular chaperone HtpG (HSP90), acts as a protein chaperone. Hsp90 exhibits diverse functions, such as helping other proteins fold correctly, maintaining the stability of these proteins under cellular stress, and helping degrade damaged or misfolded proteins in cells. These functions make Hsp90 a key control molecule in the maintenance of protein homeostasis in cells [22]. At the same time, Hsp70 and Hsp90 also participate in the oestrogen signalling pathway, and in the CK -vs.- F group, the genes involved in the synthesis of these two proteins showed an upward trend, while most of the genes in the F -vs.- AF group showed a downward trend. In addition, the MAPK signalling pathway plays an important role in regulating extracellular signal transduction, growth and differentiation in fungi [23], and the effect of MAPK on virulence has been reported in previous studies of fungal diseases [24]. The HSP70 gene involved in this pathway is upregulated in the CK -vs.- F group. Some genes in the F -vs.- AF group showed a downward trend.

## Materials and Methods

### Materials

‘HN48’ is a high-protein soybean variety. The fat content was 19.5%, and the protein content was 45.3%. The Soxhlet extraction method was used to determine the fat content, and the Kjeldahl method was used to determine the protein content [25]. The experiment was carried out at the experimental station of the Research Institute of Sugar Industry, Harbin Institute of Technology. Sterilized soil from continuous cropping of soybean was selected for the pot experiment. *F. oxysporum* was provided by the Key Laboratory of Microbiology, Heilongjiang University. *F. mosseae* was screened by our research group and stored at the Wuhan Institute of Microbiology, China. The preservation number was CGMCC no. 3013.

### Sample treatment, root rot incidence and soybean quality determination

The experiment was carried out in pots, every pot is plastic bucket and mesures 50× 50× 60 cm. Soybean seeds were wiped with alcohol, disinfected with 5% sodium hypochlorite, washed with sterile water and sown in sterilized soil from continuous cropping of soybean. Three treatments were set up: control group (CK); soybean planted in continuously cropped soil, and inoculated with *F. oxysporum* spore suspension (the concentration of spores was 1×10^7^ CFU/mL) at 44 d after sowing [26] (F) (*F. oxysporum* was activated by PDA medium) [27]; and soybean planted in continuously cropped soil inoculated at 44 d after sowing and evenly mixed with *F. mosseae* (AF). Samples were obtained every 7 days after *F. oxysporum* inoculation, and the incidence of root disease was counted[28]. During the period of high incidence of root rot (79 d), soybean roots were taken from each treatment and subjected to transcriptomic analysis. After maturation, the 100-grain weight, crude fat content and protein content were measured for each treated grain. Each treatment had three duplicates. The correlation analysis between the incidence of root rot and soybean quality indicators were carried out using SPSS 23.0.

### Transcriptomic Analysis of *F. oxysporum* in Soybean Roots

Total RNA was extracted from the samples and amplified by PCR. The whole library was prepared. The constructed library was subjected to Illumina HiSeqTM sequencing. The obtained data were filtered as follows: 1) reads containing adapters were removed; 2) reads containing more than 10% unknown nucleotides (N) were removed; and 3) low-quality reads containing more than 50% low-quality (Q-value ≤ 20) bases were removed. High-quality clean data were thus obtained. The expression levels of the identified genes were determined using RSEM software[29], and box plots comparing the gene expression levels between groups were generated. Principal component analysis (PCA) was performed with the R package gmodels (http://www.r-project.org/). To identify DEGs across samples or groups, the edgeR package (http://www.r-project.org/) was used. We identified genes with a fold change ≥ 2 and a false discovery rate (FDR) <0.05 in a comparison as significant DEGs (DEGs). DEGs identified were functionally annotated and classified based on the Gene Ontology (GO) database (http://www.geneontology.org/) and Kyoto Encyclopedia of Genes and Genomes (KEGG) database (http://www.genome.jp/kegg/pathway.html).

## Data analysis

Data were analysed by analysis of variance (ANOVA), followed by Tukey’s HSD test, to determine the significance of differences between treatments.

Hierarchical clustering analysis using Pearson correlation and principal component analysis were carried out using SPSS 23.0.

## Conclusion

Root rot is a soil-borne fungal disease caused by *F. oxysporum* that greatly reduces soybean yield. Most of the studies to date have been limited to the pathogen *F. oxysporum* alone. *F. mossea*e, as a dominant strain of AMF, plays an important role in improving plant disease resistance. In this study, transcriptomic sequencing technology was used, and the following conclusions were obtained from transcriptomic data analysis. In the AF -vs.- F group, 56 genes were upregulated, and 35 genes were downregulated. The upregulated genes mainly encoded ABC transporter-related proteins, while downregulated genes mainly encoded chitin-binding domain proteins, key enzymes in metabolic pathways, enzymes for hydrolysis of cellulose and hemicellulose, and cytoskeleton components. DEGs were enriched in antigen processing and presentation, carbon fixation in photosynthetic organisms, glycolysis/gluconeogenesis, the MAPK signalling pathway, protein processing in the endoplasmic reticulum, RNA degradation and other pathways related to fungal virulence. Thus, inoculation with *F. mosseae* can promote the growth and development of soybean and improve the disease resistance of soybean.

In conclusion, inoculation with *F. mosseae* promotes changes in gene transcription in *F. oxysporum*. The incidence of soybean root rot was significantly reduced, and the quality of soybean was significantly improved after inoculation with *F. mosseae*. Therefore, *F. mosseae* can enhance disease resistance and promote plant growth and development.

## Data accessibility

The database supporting the conclusions of this article are included. The mass spectrometry RNA data are available in the NCBI database, and the data set identifier is SRA SRP240183.

## Author Contributions

B.-Y.C. and N.G. conceived, designed and performed the experiments; X.-Q.Z. analyzed the data and contributed analysis tools; L. B. editing the manuscript; X.-Q.Z. wrote the paper. All authors read and approved the final manuscript.

## Acknowledgements

This work was supported by a grant from the National Natural Science Foundation of China [grant number 31570487, 31972502].

## Supporting information

**S1 Table**. GO analysis of DEGs in the CK-vs.-F comparison.

**S2 Table**. GO analysis of DEGs in the F-vs.-AF comparison.

**S1 Table**. KO analysis of DEPs the CK-vs.-F comparison

**S2 Table**. KO analysis of DEPs the F-vs.-AF comparison

